# Simulating Offshore Oilfield Conditions: Insights into Microbiologically Influenced Corrosion from a Dual Anaerobic Biofilm Reactor

**DOI:** 10.1101/2024.11.11.622982

**Authors:** Liam Jones, Maria Salta, Torben Lund Skovhus, Kathryn Thomas, Timothy Illson, Julian Wharton, Jeremy Webb

## Abstract

Oilfield systems are a multifaceted ecological niche which consistently experience microbiologically influenced corrosion. However, simulating environmental conditions of an offshore system within the laboratory is notoriously difficult. A novel dual anaerobic biofilm reactor protocol allowed a complex mixed-species marine biofilm to be studied. Interestingly, electroactive corrosive bacteria and fermentative electroactive bacteria growth was supported within the biofilm microenvironment. Critically, the biotic condition exhibited pits with a greater average area which is characteristic of microbiologically influenced corrosion. This research seeks to bridge the gap between experimental and real-world scenarios, ultimately enhancing the reliability of biofilm management strategies in the industry.

**Importance:** It is becoming more widely understood that any investigation of microbiologically influenced corrosion requires a multidisciplinary focus on multiple lines of evidence. While there are numerous standards available to guide specific types of testing, there are none that focus on integrating biofilm testing. By developing a novel dual anaerobic reactor model to study biofilms, insights into the different abiotic and biotic corrosion mechanisms under relevant environmental conditions can be gained. Using multiple lines of evidence to gain a holistic understanding, more sustainable prevention and mitigation strategies can be designed. To our knowledge, this is the first time all these metrics have been combined in one unified approach. The overall aim for this paper is to explore recent advances in biofilm testing and corrosion research, to provide recommendations for future standards being drafted. However, it is important to note that this article itself is not intending to serve as a standard.

## 1 Introduction

The formation and behaviour of biofilm communities in oil and gas systems, particularly those involving carbon steel (CS) surfaces in contact with produced water (PW), is a critical area of study due to its implications for material and infrastructure integrity. The presence of certain types of microorganisms can contribute to infrastructure complications, such as corrosion, souring, and biofouling [1, 2]. While some metal loss is expected and accounted for during the design of the infrastructure, higher rates of corrosion, unless detected and mitigated early, can necessitate costly repairs or replacements [3]. PW is one of the most common sampling sources within oilfield systems but is often characterised by low biomass and diversity [4]. Moreover, environmental samples are notoriously difficult to culture using standard laboratory techniques. This is because PW typically contains a diverse and complex mixture of organic compounds, including hydrocarbons and organic acids. Whereas, PW does not contain rich carbon sources, including amino acids, peptides, and sugars. Thus, simulating the environmental conditions of an offshore system within the laboratory, whilst supporting growth of environmental cultures, can be challenging. Consequently, laboratory studies on biofilms have typically been conducted under simplified conditions. However, there is a need to replicate the complex environmental parameters of offshore systems more closely to ensure the relevance of findings and any mitigation strategies employed.

Microbiologically influenced corrosion (MIC) is notoriously difficult to detect and monitor within industry. Despite the advent of molecular tools and improved microbial monitoring strategies for oil and gas operations, specific underlying MIC mechanisms in pipelines remain largely enigmatic [3]. Consequently, MIC is a particularly difficult corrosion mechanism to manage. Oilfield systems contain multiphase fluids, including crude oil, gas, as well as PW [5]. This multifaceted ecological niche consists of many undetermined thermophilic and mesophilic archaea/bacteria [5]. These uncharacterised microorganisms may metabolise organic and inorganic compounds in the crude oil and metal pipelines under extreme environmental conditions [6, 7]. Moreover, these microbial communities have the potential to change the redox chemistry within oilfield systems [8, 9]. Furthermore, PW may also contain high concentration of minerals, such as Cl^−^ and 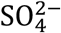, which may influence the community dynamics within the biofilm. This will impact initial biofilm formation and influence the predominant microbial metabolisms present. Therefore, any MIC processes at the interface of the biofilm and the metal surface will be affected [9, 10, 11].

The most common corrosion mechanisms reported in oilfield systems are linked to H_2_S and/or CO_2_ acid gases. These gases readily dissolve into PW, resulting in aqueous corrosive environments [3]. However, these are abiotic corrosion mechanisms. Abiotic corrosion mechanisms, such as acid gas corrosion, can be modelled *in silico* [12] or alternatively, they can be simulated in laboratory reactors [13] to understand the rates of metal loss expected under specific field conditions. This is critical for industry, as field conditions such as flow rate, temperature, water chemistry, and partial pressure of acid gases will all impact reported corrosion rates (*CR*s) [3]. Additionally, laboratory testing of abiotic corrosion mechanisms provides a robust approach for testing the efficacy of different mitigation strategies. Conversely, biotic corrosion mechanisms have not been adequately modelled within the laboratory for mixed-species biofilms from an industrial perspective. Similarly, the efficacy of biocides, applied to control MIC, have also not been adequately modelled. Thus, the development of such a model protocol is important for the effective management of MIC mechanisms. Currently, field-based detection and monitoring is primarily employed for the effective management of MIC corrosion mechanisms [3]. Microbiological assessment is routinely performed to detect the presence of MIC causative microorganisms and to evaluate the effectiveness of biocide treatments used to mitigate against MIC [14]. Though, by the time MIC is detected and diagnosed as the root cause, it may already be too late to effectively mitigate due to the recalcitrant nature of biofilms.

It is well documented that it is sessile microorganisms within biofilms that attach to metal surfaces which lead to electrochemical MIC processes. Additionally, it is widely accepted that there are distinct phylogenetic groups such as iron reducing bacteria (IRB), sulphate reducing bacteria (SRB), and acid producing bacteria (APB), which actively participate in MIC [5, 15, 16, 17]. However, long-term studies on the influence of mixed-species biofilms on MIC in oilfield systems are limited. Most laboratory studies on MIC are typically short- to medium-term, often focusing on specific aspects of the corrosion process, such as biofilm formation, *CR*, or the identification of corrosive microbial species. While these studies provide valuable insights, they may not fully capture the long-term dynamics and cumulative effects of MIC, particularly under conditions that closely mimic real-world environments. Recent studies from Elumalai *et al*. found that crude oil reservoirs were dominated by Proteobacteria, Actinobacteria, and Firmicutes classes [5, 18]. Proteobacteria biofilms have been associated with various types of corrosion, with *Pseudomonas* sp. being shown to form a thin biofilm with corrosion deposits and causing the reduction of Fe^3+^ to Fe^2+^ on the metal surface [19, 20]. Moreover, Proteobacteria species have been reported to consume hydrocarbons as a nutrient source and have the proficiency to degrade aromatic hydrocarbons for their metabolic pathways [21].

Long-term studies are challenging to conduct due to the time, resources, and complexities involved in replicating the dynamic and multifaceted conditions found in oil and gas systems. However, the need for such studies is increasingly recognised, as they can provide more comprehensive data on how biofilms evolve over time, how microbial communities interact with materials, and how these interactions contribute to long-term corrosion processes. Understanding these factors is crucial for developing effective mitigation strategies to protect infrastructure in the oil and gas industry.

This study investigates the impact of these more realistic environmental conditions on natural mixed-species biofilm communities, aiming to provide deeper insights into their development, community dynamics, and potential to induce MIC within a novel dual bioreactor protocol that simulates offshore oil and gas environments. By aligning laboratory conditions with those encountered *in situ*, this research seeks to bridge the gap between experimental and real-world scenarios, ultimately enhancing the reliability of biofilm management strategies in the industry. Importantly, utilising MLOE [22], the protocol incorporates a multi-disciplinary approach to gain a holistic understanding of biofilms and MIC.

## 2 Materials and Methods

### Test Conditions

Two anaerobic CDC (Centers for Disease Control) biofilm reactors (Biosurface Technologies Corporation) were used: an abiotic control reactor and a biotic test reactor (key dimensions: 22 cm reactor height and 12 cm internal diameter; 21 cm coupon holder rod; 1.27 cm coupon diameter). Sterile carbon steel coupons were fixed in reactors and exposed to two separate conditions for 28 days. Anaerobic conditions were maintained throughout the test by initially sparging the system with nitrogen gas (oxygen free nitrogen) (BOC Nitrogen (Oxygen Free), 44-W) over an initial three-day batch phase. Anaerobic conditions, considered to be hypoxic or low O_2_, are characterised as a system with low concentrations ranging between 1 and 30% saturation. Strict obligate anaerobes will not survive if there is more than half a percent O_2_ in the environment, while moderate obligate anaerobes can still grow in a 2 to 8% O_2_ environment [23]. PW collected from an offshore platform (1st stage separator: 50 °C) was used as the test medium. The bulk-produced water, used in the reactors, was prepared by filter sterilisation prior to use to ensure sterility, using a 0.2 µm Vivaflow® TFF Cassette, PES (Sartorius). The test media composition can be seen in Supplementary Table S1. Resazurin was added as a redox indicator, as it is colourless under oxygen free conditions but changes to a pink colour in an oxygen-containing environment. Agitation of the reactor baffles was set to 50 rpm to maintain a homogeneous solution. The reactor temperature was set to 40°C, to better simulate the environmental condition of the offshore system. Prior to inoculating the biotic reactor, a three-day pre-culture was prepared in a blue-cap flask (50 mL), consisting of 10% marine sediment with the remainder filter sterilised PW media. The pre-culture PW was prepared by adding 1 g/L of yeast extract to filtered produced water and autoclaved. It was acknowledged, for the pre-culture, that yeast extract contains redox mediators that may adsorb onto the electrode surfaces and chelate metal ions and the test matrix was designed to highlight any significant interference [24]. The biotic reactor was inoculated using a sterile syringe, where 10% of the working reactor volume (35 mL) was added as the inoculum. Initial adenosine triphosphate (ATP) measurements were taken from the pre-culture and long-term frozen stocks were prepared using 20% glycerol. Supplementary Figure 2(a) shows a schematic of the full experimental setup, with Supplementary Figure 2(b) illustrating the three-electrode cell setup within each anaerobic CDC biofilm reactor. Both reactors were operated in batch mode for the first three days to allow settlement and to facilitate biofilm formation in the biotic reactor. After this period, the reactors were switched to continuous flow of fresh media at a rate of 0.2 mL min^−1^, which replaced about 50% of the 600 mL total volume daily (288 mL day^-1^).

### Microbial consortia

The sheltered zone littoral sediment microbial consortia were collected at a depth between 10 – 15 cm below the sediment surface during low tide from Langstone Harbour, United Kingdom (50°50’11.9”N 0°58’47.5”W). The coastal/estuarine marine sediment (very fine and cohesive mud and silt deposits) was selected to sample microorganisms living under low oxygen conditions. The sediment samples were added to 500 mL of the PW medium and stored at 37°C in an anaerobic chamber to maximize the recovery of the diverse microbial populations. Mesophilic bacteria can survive and grow in temperatures between 10 – 50°C. Thus, a tropic strategy to promote cell growth and viability was employed to maximise microbial recovery. The anaerobic chamber gas mixture consisted of 85% N_2_, 10% CO_2_ and 5% H_2_ (BOC Anaerobic Growth Mix, 290563-L). Long-term storage of sediment samples and microbial consortia was employed to create frozen stocks at –80ºC.

### Carbon steel coupon preparation

UNS G10180 (AISI 1018) carbon steel disc coupons (Biosurface Technologies – RD128 CS), with dimensions of 12.7 mm diameter × 3.8 mm thickness were used as-received (AR) (*R*_a_ = 1.31906 ± 0.33859). The surface profiles and weights for all coupon samples were assessed prior to starting the experiment, on Day 0, for surface profilometry and gravimetric analysis to be performed at the completion of the experiment after Day 28. Three-dimensional (3D) surface profiles were taken using a 3D optical profilometer (Alicona imaging infinite focus microscope IFM G4 3.5). A Mettler AT201 was used to take five measurements of the initial weights of all coupons.

### Experimental setup

Before autoclaving, the two anaerobic CDC biofilm reactors were cleaned with detergent and allowed to dry. The empty reactors with attached tubing were placed in autoclavable bags; all tube openings and air filters (Millex, 0.2 µm) were covered in aluminium foil, with tube openings clamp shut. The empty assembled reactors were autoclaved for 15 mins at 121°C. After cooling the reactors were transferred into a sterilized microbiological safety cabinet, along with all rods, carbon steel test coupons, as well as any sensors and electrodes. Working electrode rods were prepared in advance. For each working electrode rod, wires were soldered to each coupon separately. The coupon face with the soldered wire was then covered with a lacquer solution (Polishing Shop, Type 45 Stop Off Lacquer) and allowed to dry. To assemble the reactors, all rods with coupons were submerged in 99% ethanol for at least 10 s, then inserted into the autoclaved reactors. Any sensors or electrodes used in place of a rod were also inserted, after also being sterilized with 99% ethanol for at least 10 s. The medium bottles and all tubing were connected in a microbiological safety cabinet. Once both reactors were fully assembled, they were transferred to the working area, with access to a N_2_ gas supply. The tubing was evenly split into each reactor to equalize the pressure gradient caused by the peristaltic pump (Matson Marlow 300 series).

### Sulphide analysis

Sulphide concentrations were monitored daily in each reactor using a Unisense, SULF-50 sulphide microsensor (50 μm diameter) and amplifier (Unisense, H_2_S UNIAMP). The microsensor measures the partial pressure of H_2_S gas, and the total concentration is a function of pH and temperature. The microsensor limit of detection is 0.3 µM, with a range from 0-300 µM sulphide in water. Calibration utilised the H_2_S and SULF sensor calibration kit (Unisense, CALKIT-H_2_S). Due to the nature of the experimental setup, it was not possible to calibrate the microsensors during the experiment. However, calibrations were performed both prior to starting the experiment and once the experiment had finished to confirm that the sensors were still calibrated. The SensorTrace Suite software was used to collect the sulphide microsensor data. The sensor has a higher signal for zero right after it has been connected to the amplifier, thus each microsensor collected readings for five minutes (approximately 300 data points) on each day. This was to allow the sensor to stabilise.

### Surface profilometry and visual inspection

Corrosion products and biofilms were removed from the surface using the cleaning protocol as described previously for the gravimetric analysis. Three-dimensional (3D) profiling of the carbon steel surfaces was reconstructed using an Alicona imaging infinite focus microscope IFM G4 3.5. The images allowed assessment of changes in surface roughness compared to the surface profiles obtained prior to testing. Additionally, ImageJ/Fiji was used for the quantitative determination of pit depth, width, height, percentage area, and to assess pit rate (*PR*) and pit density (*PD*). This analysis was performed on thirty total locations on six AR coupons (five locations each). The method involved applying a colour threshold to depths greater than 5 µm. Then, the images were converted to a binary mask. Next, measurement parameters were selected for areas greater than 650 µm^2^. Finally, the images were analysed to display counts, area, and average size of pits. The pit parameters were adapted from ASTM G48-11 [25]. For pit rate analysis, the deepest pits from each image were captured using the Alicona. Pit rates were calculated using the formula described in NACE SP0775-2023 [26].

### Gravimetric analysis

Corrosion products and biofilms were removed following the ASTM G1-03 standard with a 15% inhibited hydrochloric acid described in NACE SP0775-2023 [26, 27]. A stock solution was made of 37.5% HCl (Merck, Suprapur, 1.00318.0500) to which 10 g/L of 1,3-di-n-butyl-2 thiourea (DBT) (Merck, 8.20423.0250) was added. Immediately prior to use, the stock solution was diluted by slowly adding a measured volume of stock solution to an equal volume of deionised water with stirring. A Mettler AT201 was used to take five measurements of all coupons. Corrosion rates were determined by the gravimetric technique that considers the weight loss and surface area of the metal samples described in NACE SP0775-2023 [26].

### Electrochemical analysis

Electrochemical measurements were performed using a Gamry Instruments potentiostat (Ref 600 Plus). The electrochemical behaviours of the carbon steel coupons were evaluated using a three- electrode system consisting of a UNS G10180 coupon as the working electrode, graphite rod (Alfa Aesar, 99.9995%, 6.15mm diameter, 152mm long) as the counter electrode, and a silver/silver chloride (Ag/AgCl, 3.5 M KCl) reference electrode (Sentek, (AgCl) Double junction Reference Electrode). On day 1, after the test reactor was inoculated, both reactors were left for at least 1 h prior to performing any electrochemical measurements. Open-circuit potentials (OCP) were recorded for each coupon on day 1 prior to measuring linear polarization resistance (LPR) and electrochemical impedance spectroscopy (EIS). LPR and EIS were measured daily for each sample. LPR measurements were performed from ±10 mV with respect to *E*_OCP_ using a scan rate of 0.167 mV s^-1^. EIS measurements were performed at OCP with an applied 10 mV_rms_ sinusoidal potential signal with a frequency range of 10^−2^ to 10^5^ Hz. Potentiodynamic polarization measurements were performed at the end of the experiment on day 28 for each coupon from –0.200 mV to +0.200 V using the scan rate of 0.5 mV s^-1^. Standard procedures were followed when selecting an equivalent circuit best-fit using the Gamry Echem Analyst software: (*i*) the chi-squared (*χ*^2^) error was suitably minimized (*χ*^2^ ≤ 10^−4^) and (*ii*) the errors associated with each element were ranged between 0 % and 5 %.

### Confocal laser scanning microscopy and post-image analysis

The distribution of live and dead cells within biofilms was studied using confocal laser scanning microscopy (CLSM). Coupons were gently rinsed with sterile anaerobic PBS, with the following composition: NaCl 8g, KCl 0.2 g, Na_2_HPO_4_ 1.44g, KH_2_PO_4_ 0.245 g, deionised water 1 L. and stained using the FilmTracer Live/Dead biofilm viability kit (Invitrogen) according to the manufacturer’s instructions. Before imaging with a Leica SP8 confocal microscope, coupons were rinsed with sterile deionized water to remove the excess of dyes and fixed using mowiol. Mowiol had the following composition: 2.4 g Mowiol, 6 mL deionized water, 12 mL 0.2 M Tris (pH 8.5), 0.01 g sodium azide, and 6 g glycerol. Images were obtained with a 63× magnification and glycerol immersion. The dyes used stained live cells with a green-fluorescent colour (SYTO 9) and dead cells with a red colour (propidium iodide). The *z*-stacked images were analysed using Imaris software (Oxford Instruments).

### Microbial community analysis

After 28 days, six AR coupons were gently rinsed with PBS and then placed in a Falcon tube containing 10 mL of PW solution. Long-term frozen stocks were prepared using 20% glycerol for the bulk fluid, AR biofilm, and P biofilm samples from the biotic reactor. The sediment, three-day pre- culture, day 28 bulk fluid, and AR biofilm frozen stocks were sent in triplicate for DNA extraction and 16S rRNA amplicon sequencing. Library preparation and sequencing were performed for the V3 and V4 regions of the 16S rRNA gene targeting both bacteria and archaea. The microbiome analysis pipeline along with DNA extraction was performed by Eurofins Genomics LLC. Taxonomic classification method using Kraken2 (v 2.1.1). Bioinformatics and data analysis was performed using the Qiime2 (version 2023.5) software. To visualize the multivariate dispersion of the community composition, a principal component analysis (PCA) analysis was conducted employing GraphPad (version 10.0.2).

### ATP assay

The ATP concentration in both the abiotic and biotic reactors was determined by luminescence after reaction with luciferin-luciferase using the BacTiter-Glo™ Microbial Cell Viability Assay kit (Promega). The assay provides a method for determining the number of viable microbial cells in culture based on quantitation of the ATP present. ATP is the energy source of all living cells and is involved in many vital biochemical reactions. When cells die, they stop synthesizing ATP and the existing ATP pool is quickly degraded. Higher ATP concentration indicates higher number of living cells. All assays were performed according to the manufacturer’s instructions. Six AR coupons were gently rinsed with PBS and then immersed in a Falcon tube containing 10 mL of PW solution. Any cells were detached from the metal coupons using a cell scraper (Biologix). Both planktonic and sessile samples were processed with the BacTiter-Glo™ Microbial Cell Viability Assay kit, which measures ATP from as few as 10 microbial cells. The ATP concentrations were determined by measuring luminescence with a Clariostar Plus Multimode Microplate Reader (BMG Labtech). Planktonic cells in each reactor were determined following the same method described before; in this case, 10 mL of the bulk test solution was processed with the BacTiter-Glo™ Microbial Cell Viability Assay kit. Negative controls of PBS, deionised water and PW solution were used to indicate no ATP activity.

### Corrosion product analysis

Analysis of the corrosion products and biofilms were performed by SEM-EDS and Raman microspectroscopy. For SEM, all images and Energy dispersive X-ray spectroscopy (EDS) measurements were taken using a Hitachi S-3400N II SEM and attached energy-dispersive X-ray spectrometer (Oxford Instruments). Imaging was performed at approximately 15 kV with a working distance of 10 mm at various magnifications (71×,1000×,and 3000×). EDS analysis used the same parameters and magnifications. For each sample analysed areas were scanned at randomly distributed areas over the sample surface (n=15). EDS data was analysed using AZTEC software before being compiled in Microsoft Excel. Raman microspectroscopy experiments were conducted using a Renishaw InVia Raman microscope (Renishaw, UK), with a Leica DM 2500-M bright field microscope and an automated 100 nm-encoded XYZ stage. The samples were excited using a 532 nm laser directed through a Nikon 50× long working distance air objective (NA = XX). Raman-scattered signals were separated from the laser illumination at 532 nm using a Rayleigh edge filter, and a diffraction grating (532nm: 1800 L/mm) dispersed the Raman-scattered light onto a Peltier-cooled CCD (1024 pixels × 256 pixels). Calibration of the Raman shift was carried out using an internal silicon wafer using the peak at 520 cm^-1^. Spectra were acquired over two or three accumulations of between 5 and 20 s each, using laser power of up to 3 mW. Spectra were acquired from a selection of points manually determined using the brightfield imaging mode of the microscope. The spectra obtained were processed using MATLAB (MathWorks).

## 3 Results

### Visual observations

Over the initial three-day batch phase, the abiotic media was pink in appearance, and the coupons maintained the silver-grey metallic lustre of CS. After the first week, the abiotic media was beginning to become cloudier, but it was not until Day 11 that the abiotic PW became orange-pink in colouration. Similarly, over the initial three-day batch phase, the biotic coupons maintained the silver-grey appearance of the CS. However, the PW quickly became dark green/black in colouration on Day 3. After the flow of fresh PW media was started on day 4, the bulk fluid became pink in appearance. Similarly, it was not until Day 11 that the biotic PW became orange-pink in colouration. Upon dismantling of the reactors on day 28, and retrieval of the coupon rods, there was a significant difference in the coupon appearances, see Supplementary Figure S3. Additionally, the appearance of the waste from the different reactor conditions was significantly different. The abiotic waste media was orange-pink in colouration with reddish-brown granular deposits, whilst the biotic waste media was dark green/black in colouration. The abiotic surfaces had the presence of a black corrosion product across the entire coupon surfaces, with some reddish-brown granular deposits. Whereas, the biotic surfaces were only partially covered by a black corrosion product, with the presence of a heterogeneous dark green/black biofilm. The silver-grey appearance of the CS was also partially visible upon retrieval.

### Sulphide analysis

All values were detected as below zero, as such it was assumed that no H_2_S was present throughout the experiment. Alternatively, the PW poisoned the microsensors. Nonetheless, both conditions experienced similar readings. Unfortunately, *DO* concentrations could not be measured as the probe was not working. However, the pH was measured on day 28 for the media containers, waste containers and reactor systems. All values recorded were between 7.01 and 7.47.

### Carbon steel surface analysis

Supplementary Figure S4 shows the CS surfaces on day 0. Supplementary Table S5 summarises the quantitative surface roughness profiles on both day 0 and day 28. Figure 1 shows the cleaned CS surfaces after 28 days, with biofilms and corrosion products removed to reveal the morphology of the surface degradation and to facilitate corrosion assessment. Surface profilometry revealed that there were low levels of uniform or localised pitting corrosion present for both the abiotic and biotic condition. The abiotic average pit depths were 12 µm, with an average pit area of 971 µm^2^ for any classified pits. The biotic average pit depths were 7 µm, with an average pit area of 1501 µm^2^ for any classified pits. Again, for this study a pit was classified as having a depth greater than 5 µm and an area greater than 650 µm^2^ [28].

**Figure 1.**
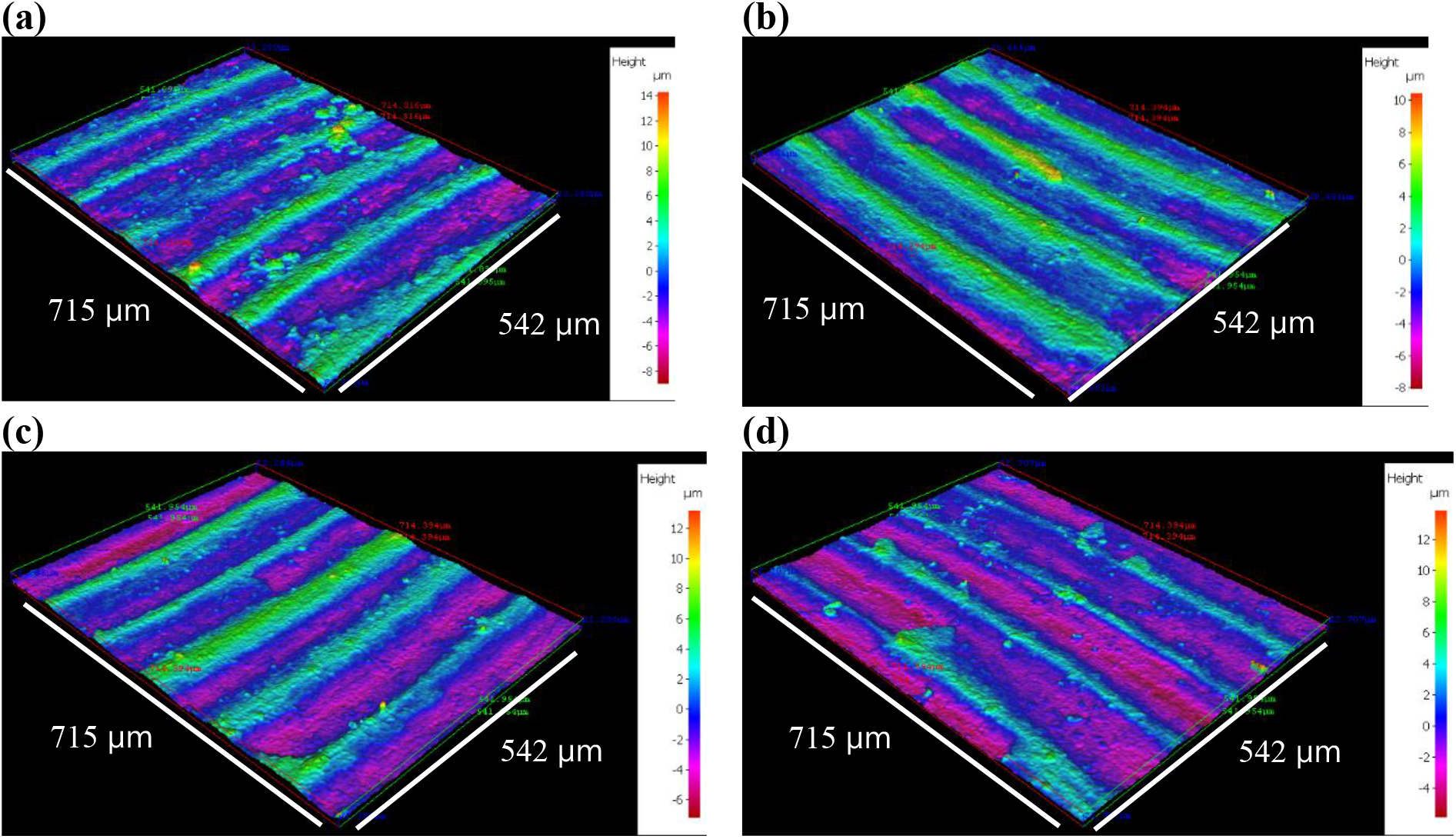
Three-dimensional optical surface profilometry of the cleaned UNS G10180 surfaces at day 28. AR coupons for: (a,b) abiotic and (c,d) biotic conditions, after exposure to anaerobic produced water media for 28 days.

Figure 2a provides an evaluation of the CS coupons *CR*. For the abiotic condition, there was a higher *CR* when compared to the biotic condition, though there was no significance. According to the NACE SP0775-2023 assessment criteria, there was a moderate *CR* (between 0.025 and 0.12 mm y^−1^) in both the abiotic and biotic reactors (Figure 2a). Whereas a moderate *PR* (0.13 – 0.20 mm y^−1^) was assessed for the abiotic condition, with a low *PR* (<0.13 mm y^−1^) for the biotic condition (Figure 2b) [29]. Further analysis of the surface profilometries in Figure 1, allowed a quantitative determination of the *PR* Figure 2b of the CS coupons. For this study, it was not possible to quantitatively determine *PD* values, due to the general absence of pitting across the coupon surfaces upon retrieval after 28 days.

**Figure 2.**
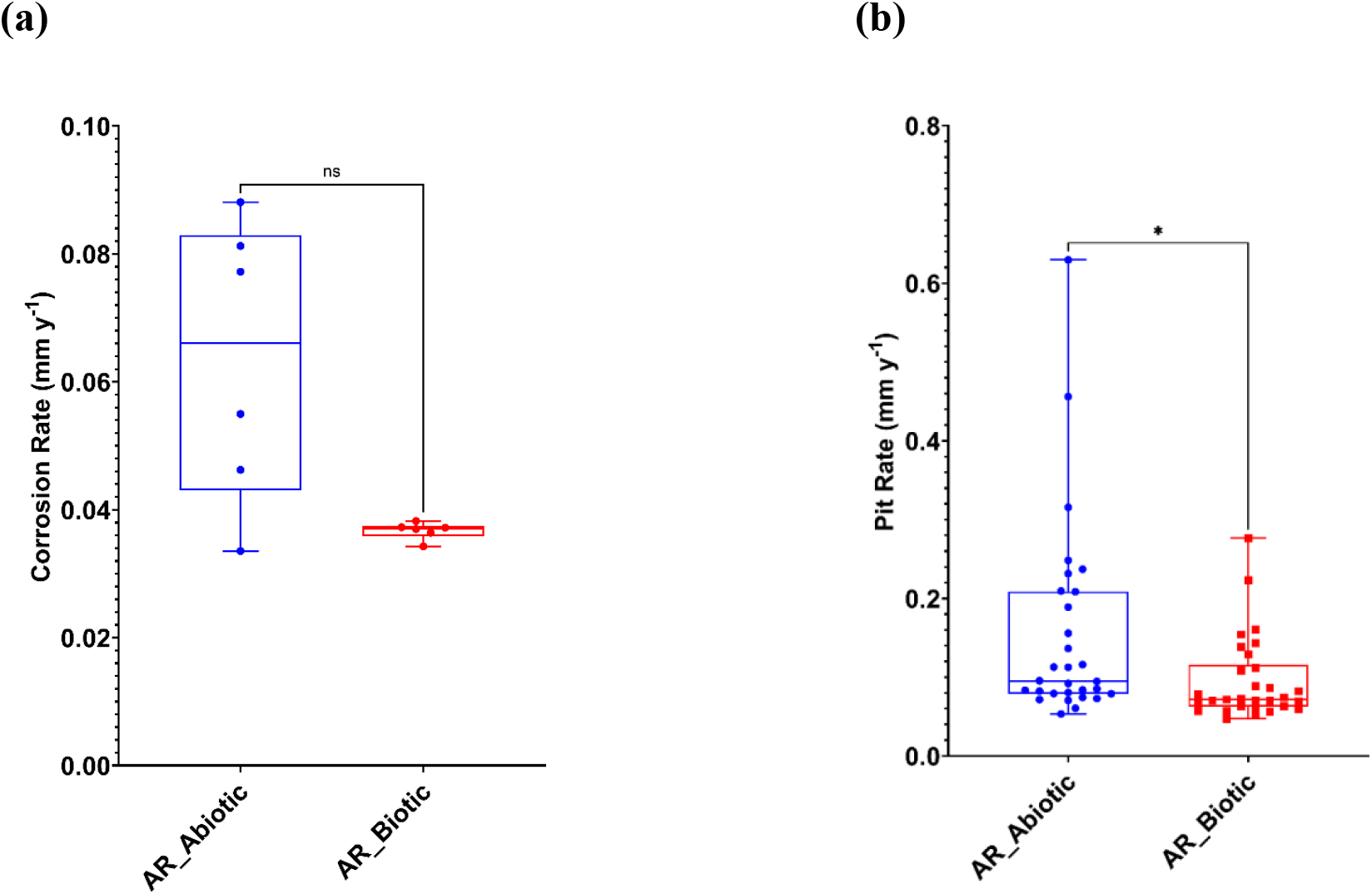
Abiotic and biotic corrosion performance after exposure to anaerobic produced water media for 28 days: (a) corrosion rate via gravimetric analysis and surface profilometry assessed and (b) pit rate (P < 0.05), for the AR coupons.

### Corrosion product analysis

Figure 3 shows SEM-EDS elemental mapping of the UNS G10180 CS surfaces for both the abiotic and biotic conditions. Quantitative SEM-EDS data collected from elemental mapping are shown in Supplementary Figure S6. The images of corrosion products and biofilms attached to the metal samples demonstrate the heterogeneity of distribution over the surface. Coupons exposed to the biotic condition were observed to exhibit greater surface coverage. The SEM-EDS elemental maps are shown in Figure 3b – d. The major elements detected in coupons exposed to all conditions were iron (Fe), sulphur (S), and oxygen (O). Corroded areas of all coupons were mainly covered by Fe and O, with heterogeneous distribution of S. A cross-sectional image of the corrosion products was not performed.

**Figure 3.**
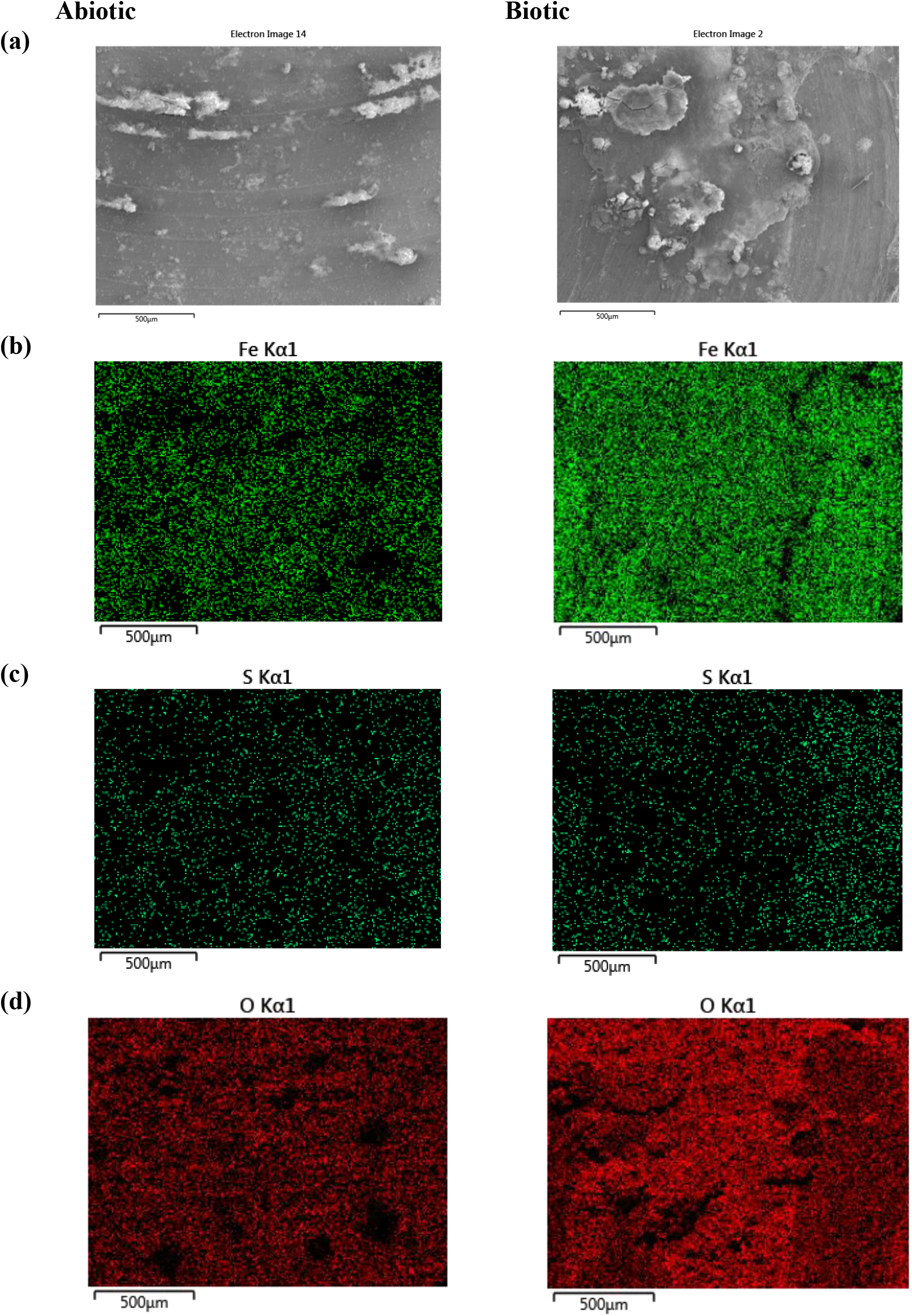
SEM-EDS elemental mapping of the UNS G10180 carbon steel, AR surfaces, after exposure to anaerobic produced water media, taken on day 28. (a) SEM image; (b) iron map; (c) sulphur map; (d) oxygen map.

Additional analysis of the corrosion products using Raman spectroscopy are shown in Figure 4. According to Raman bands of reference corrosion products in previous papers [30, 31, 32], the corrosion products are identified to be primarily mackinawite (bands 208, 282 cm^−1^) for both the abiotic and biotic condition. There were also additional bands which may be attributed to sulphur, as well as reference iron oxide compounds such as magnetite, goethite (*α* − FeO(OH)), lepidocrocite, or hematite (*α* − Fe_2_O_3_). The composition of this black compact layer was identified at mid-strong bands 250, 380, 1307 cm^−1^ associated with lepidocrocite. Additionally, bands at 298, 399, 481, 554, 675 and 1002 cm^−1^ have previously been shown to be associated with goethite, whilst bands at 222, 244, 298, 501, 615 and 1318 cm^−1^ are associated with hematite. Magnetite has previously been shown to be associated with bands at 675 and 550 cm^−1^ [30, 31, 32]. The coverage of the metal sample with a black precipitate was indicative of the successful growth of corrosion products film containing FeS compounds. The Raman spectrum of the sample is in good agreement with literature spectra attributed to mackinawite.

**Figure 4.**
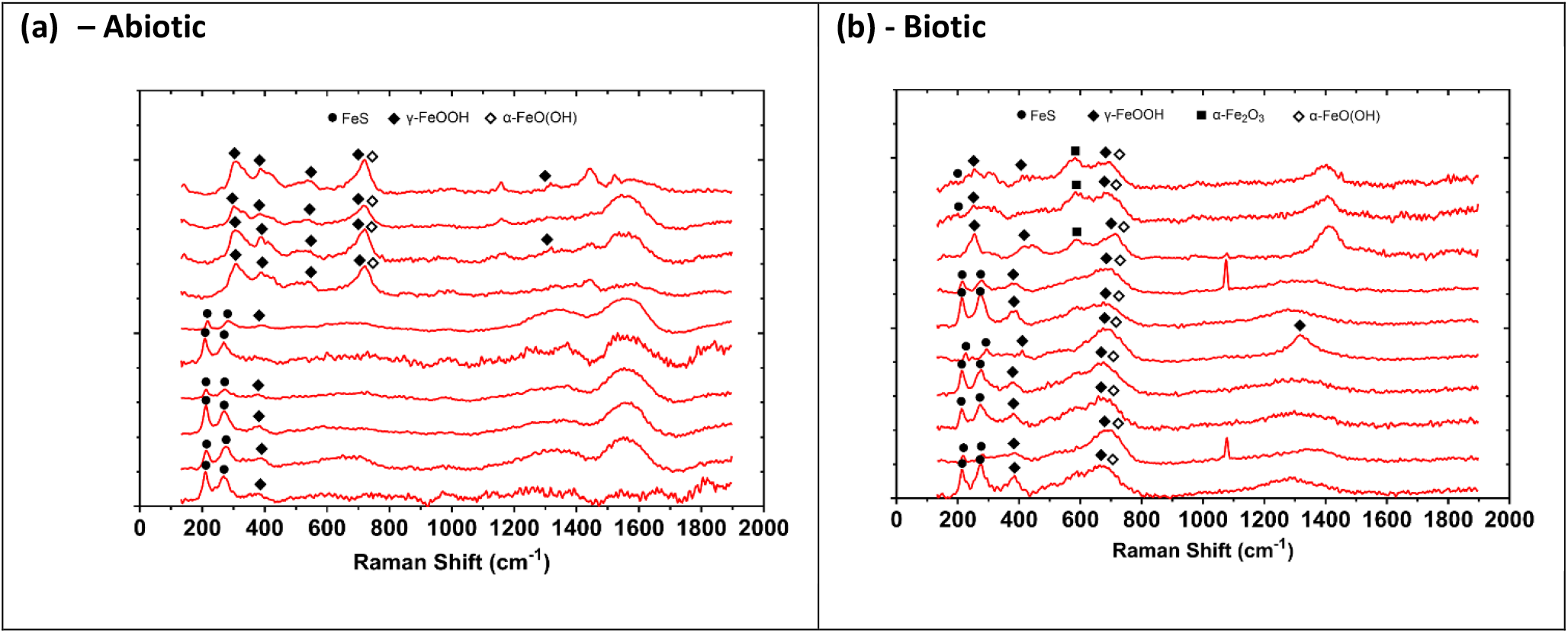
Raman spectra of the UNS G10180 carbon steel surfaces after exposure to anaerobic produced water media, taken on day 28. For the AR (a,b) abiotic and; (c, d) biotic condition.

### Electrochemical measurements

Figure 5 shows the changes in *E*_corr_ and *R*_p_ between the abiotic and biotic anaerobic PW media, for the UNS G10180 CS coupons. For the abiotic condition, Figure 5a, there was a swift increase of +0.040 V on Day 3 after the end of the batch phase. This was subsequently followed by swift decrease of -0.050 V on Day 4. This can be attributed to the flow of fresh PW. Otherwise, there was a gradual +0.090 V electronegative shift in the *E*_corr_ between days 4 and 28. This can be linked with the presence of a conditioning film (i.e., an adsorbed organic layer) and the formation of inorganic corrosion product layer. A pseudo-steady state *E*_corr_ had not been attained after 28 days. Similarly, for the biotic condition, there was a swift increase of +0.050 V on Day 3 after the end of the batch phase. This was subsequently followed by swift decrease of -0.030 V on Day 4. Then, there was a gradual electronegative shift in the *E*_corr_, until Day 24. After which, the *E*_corr_ swiftly decreased by -0.040 V. The potentials for both abiotic and biotic in the latter stages were generally similar and ranged between –0.610 V and – 0.670 V *vs*. Ag/AgCl.

**Figure 5.**
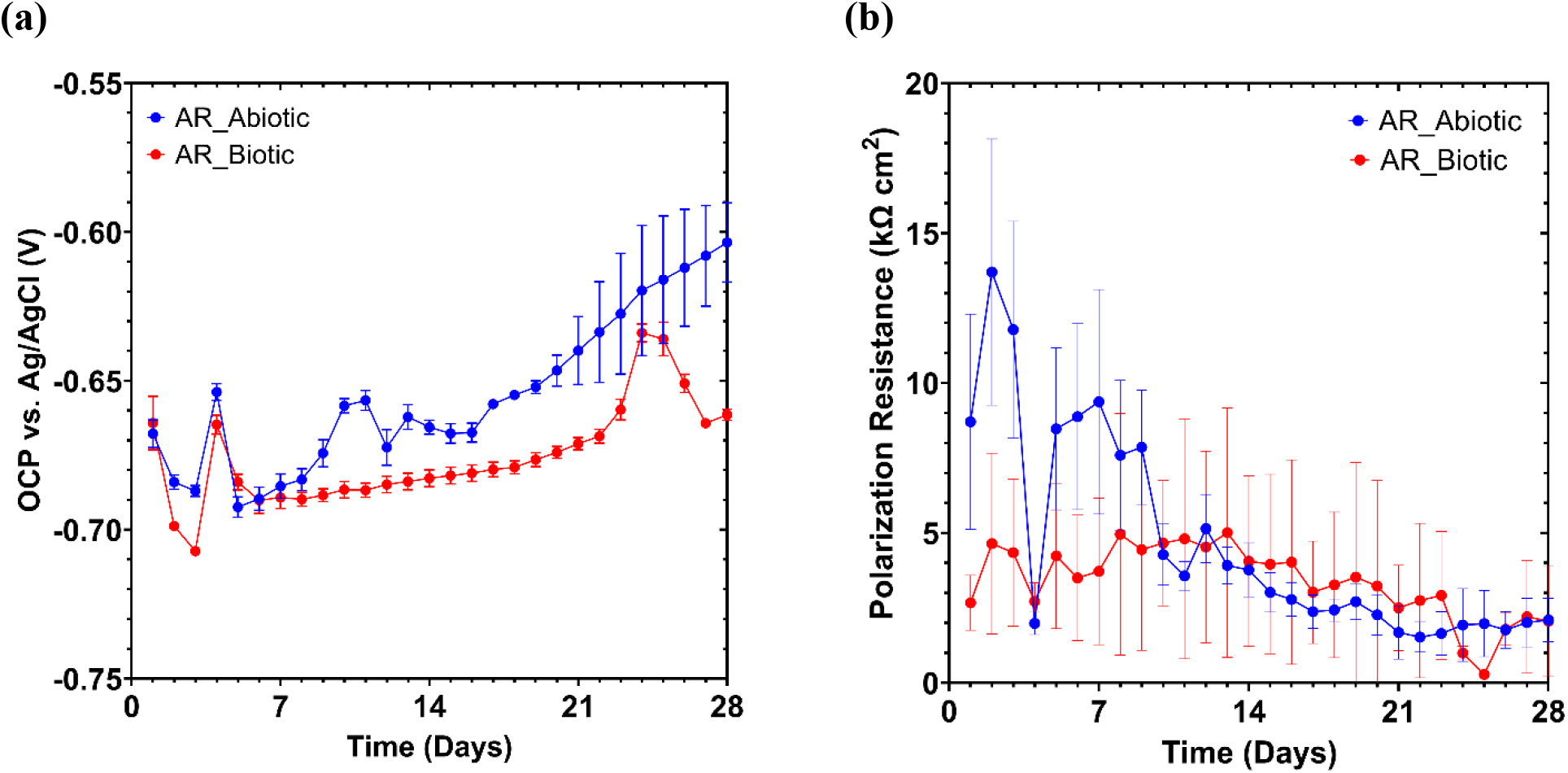
LPR data for UNS G10180 carbon steel: (a) open-circuit potentials and (b) polarisation resistance in anaerobic produced water media (abiotic and biotic conditions), for AR and P coupons (data points represent mean ± standard deviation, *n* = 6). Reactor stirrer at 50 rpm.

In Figure 5b the LPR derived *R*_p_ after Day 10 remained low at approx. 400 Ω cm^2^ for the sterile abiotic condition, indicative of a uniform corrosion across a porous corrosion film. Similarly, for the biotic condition, the *R*_p_ remained low at approx. 400 Ω cm^2^. The pioneering bacterial attachment/colonisation was difficult to detect for this study. However, biofilm formation and growth kinetics will lead inevitably to a more complex electrochemical response. Overall, there were no significant differences when comparing between the abiotic and biotic reactor environments.

Figure 6 shows the EIS data for UNS G10180 CS in the anaerobic PW media presented in three forms: Nyquist, Bode phase angle and Bode impedance modulus plots. The sterile abiotic condition on Day 1 typifies an electrochemical response for the formation of a porous interface, with diffusion of soluble electroactive species across an organic conditioning film [33] and nascent inorganic corrosion product layer. The diffusive behaviour is associated with linear features having a roughly 45° slope (a Warburg impedance response) and phase angles close to 45° in low frequency region (10^− 2^ – 10^0^ Hz), see Fig. 6a and 6c. This electrochemical response did not really change over the 28 days. Similarly, the biotic condition had a consistently uniform EIS response over the 28 day test, with only minor variation in the spectra and suggests the absence of significant detectable electrochemical changes with time. Notably, there are no discernible Nyquist semicircles (Fig 6b). Here a wider low frequency region (10^−1^ – 10^2^) is likely to be subject to a greater influence of adsorption processes, associated with the adhesion of the pioneering bacteria on a conditioning film [34, 35] and biofilm formation. The EIS spectra were fitted using an ECM shown in Supplementary Figure S7. Both the abiotic and biotic data generally had a good fit, with the quantitative fitting results shown in Supplementary Table S8.

**Figure 6.**
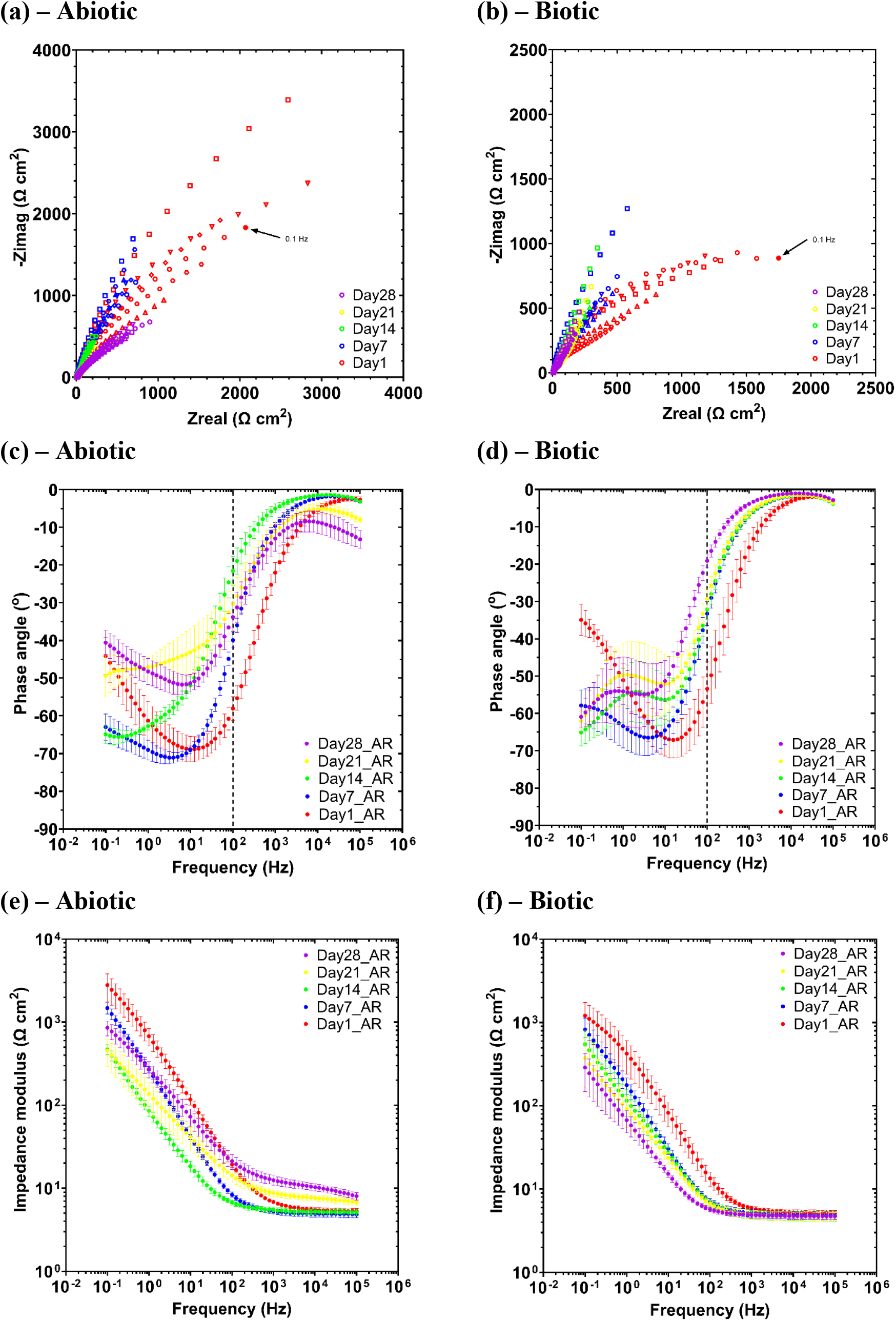
EIS data for UNS G10180 carbon steel in anaerobic produced water media at OCP: (a, b) Nyquist, (c, d) Bode phase angle (*θ vs. f*), and (e, f) Bode impedance modulus (|*Z*| *vs. f*) over 28-days. (*n* = 6). Reactor stirrer at 50 rpm.

For the abiotic control, there is a capacitive behaviour over the first week, with a diffusive behaviour over the final three weeks in the film layer (reflecting ion adsorption). Whilst there was a diffusive behaviour in the double layer, which reflects charge transfer, due to the formation of corrosion products (rust, porous oxide layer). The exponent parameter in the double layer reflects a non-ideal capacitance, which is indicative of resistive and inductive parasitics, because of a more prominent resistive component. *R*_film_ is relatively low over the 28 days, and there are no significant changes in *R*_ct_. Similarly, for the biotic condition, there was a capacitive behaviour over the first week, with a diffusive behaviour over the final three weeks in the film layer. Whereas there was a diffusive behaviour over the initial two week, followed by a capacitive behaviour over the final two weeks in the double layer. This capacitive behaviour is attributed to the biofilm. There are no significant changes in the *R*_ct_ in the double layer over time. The exponent parameter for the film layer is greater than 0.8 only on day 28, which indicates a non-ideal capacitance response. This is true for the final two weeks in the double layer. The ECM and EIS both have general agreement with the LPR data.

Supplementary Figure S9 shows the potentiodynamic polarisation curves for UNS G10180 CS for the abiotic and biotic reactors in anaerobic PW media after 28 days. Supplementary Table S10 shows the corrosion parameters obtained from the polarisation curves. From the Tafel slopes, there is a similar cathodic behaviour (reduction) when comparing the abiotic and biotic conditions, which is linked to the predominant HER under anaerobic conditions. Conversely, the anodic Tafel slopes (oxidation) are greater, demonstrating almost limiting current densities, in the abiotic compared to the biotic media. Overall, the abiotic condition had a higher *j*_corr_ compared to the biotic condition. This is consistent with a more uniform corrosion morphology. Similarly, the sterile abiotic condition had a more electropositive *E*_corr_ when compared to the biotic condition. The polarisation results corroborate the LPR and EIS data.

### Biofilm characterisation

CLSM with differentiation of live and dead biofilm cells was performed and can be found in the supplementary material Supplementary Figure S11. The heterogeneous biofilm distribution over the surface of the CS coupons did not allow measurements of the maximum biofilm thickness. Therefore, the thickness of biofilms was not determined. From the images captured, there was a live/dead cell ratio of approximately 87% live to 13% dead.

Active microorganism evaluation of the environmental marine sediment, the initial and final biotic PW media planktonic samples (Day 0 and Day 28), and the biotic AR biofilm, was undertaken via 16S rRNA amplicon sequencing with two target region, V3-4 for bacteria and archaea. A total of 2,293,909 high-quality sequences were obtained after bioinformatics processing of the raw reads. From these, 97.7% was classified for the sediment sample with 99.99% classified for the Day 0 planktonic sample and 100% classified for the Day 28 planktonic sample and AR biofilm sample. These sequences were taxonomically classified into microbial genera. The top 25 microbial genera are presented in Supplementary Table S12 in the supplemental material. Figure 7 summarises the sequencing data, showing a PCA (a) and a stacked bar plot (b) illustrating the relative abundances for the top 25 genera. Molecular identification of the microorganisms showed that the initial sediment sample had a very diverse microbial composition. Most genus had low relative abundances less than 2%. The dominant genera included *Sulfurovum, Candidatus Prometheoarchaeum, Desulfosarcina, Desulfuromonas* and *Thiohalobacter*. Interestingly, there were relatively high numbers of archaea in the sediment sample compared to the other samples.

**Figure 9.**
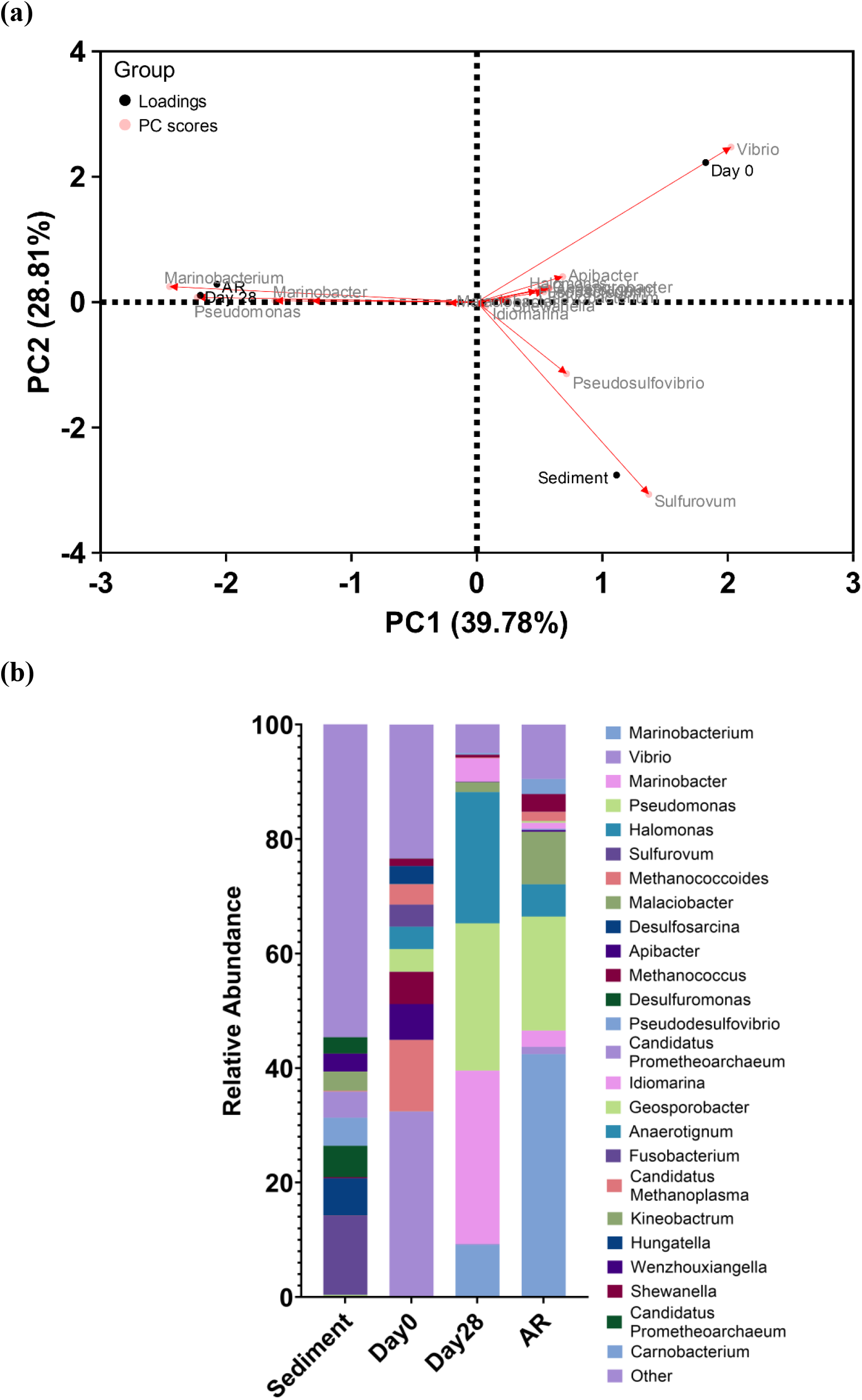
Principal Component Analysis biplot (a); Microbial community. The results show the mean relative abundances of microbial communities classified at the genus level, for the top 25 genera, from 16S rRNA amplicon sequencing (b); for environmental marine sediment, Day 0, and Day 28 planktonic samples, and AR biofilms, after exposure to anaerobic produced water media for 28 days.

The sediment sample had negative Spearman correlation coefficients (Supplementary Figure S13) with the other samples which was attributed to changes in conditions such as temperature and media composition from the natural marine environment. There was much less diversity in the Day 0 sample, with *Sulfurovum, Candidatus Prometheoarchaeum* and *Thiohalobacter* all exhibiting negligible relative abundances. Whilst genera from *Vibrio, Methanococcoides, Apibacter*, and *Methanococcus* made-up approximately 55% of the relative abundance. Again, the Day 0 planktonic sample had negative Spearman correlation coefficients with the other samples. After 28 Days, there was a significant shift in the microbial composition, with substantially lower abundances of methanogenic species. Conversely, there was a significant increase in Proteobacteria species. *Marinobacter, Pseudomonas*, and *Halomonas* were the dominant genera making up approximately 80% of the relative abundance. The Day 28 planktonic sample had a Spearman correlation coefficient of -0.43 with the sediment sample, -0.38 with the Day 0 planktonic sample and 0.89 with the AR biofilm sample. The relative abundances of *Vibrio* decreased to approximately 1%, with the relative abundances of *Methanococcoides, Apibacter*, and *Methanococcus* also decreasing to negligible values in the AR biofilm. Moreover, *Sulfurovum, Candidatus Prometheoarchaeum, Desulfosarcina, Desulfuromonas* and *Thiohalobacter* which were the dominant genera from the sediment sample all had negligible relative abundances in the biofilm sample. The dominant genera included *Marinobacterium, Pseudomonas*, and *Malaciobacter* making up approximately 70% of the relative abundance. There were no methanogenic archaea in the biofilm sample.

The microbial activity was determined by the ATP concentrations (dissolved, dATP) in the bulk fluid, see Supplementary Figure S14. The ATP assay did not measure any ATP from the biofilm sample. The biofilm may have been loosely adherent to the coupon surfaces are may have been rinsed off during sample preparation. For the biotic PW media (bulk fluid), there was a significant change (P < 0.05) in the dATP concentration when comparing Day 0 and Day 28, with dATP values of on the order of 100 pg mL^−1^. As expected, there was a significantly greater ATP concentration for the biotic compared to the abiotic condition.

## 4 Discussion

Case studies from offshore oilfield systems have consistently demonstrated that environmental factors such as temperature, salinity, O_2_ levels, and the composition of PW play a crucial role in the development of biofilms and the severity of MIC on CS surfaces [36]. These studies underscore the importance of implementing tailored corrosion management strategies that include regular monitoring, biocide application, material selection, and maintenance to mitigate the threat of MIC in offshore oil and gas operations. However, there are limitations and challenges when testing *in situ*. Offshore environments are often remote and difficult to access, making it challenging to collect samples and monitor corrosion processes over time. Environmental variability, including changes in temperature, pressure, salinity, and flow rates can further complicate the identification of specific factors driving MIC. Continuous long-term monitoring is essential to fully understand the dynamics of biofilm development and MIC in offshore systems. However, maintaining long-term studies in such harsh environments is challenging. Consequently, this leads to a reliance on shorter-term studies that may miss important trends. This can make it difficult to identify and characterise the full range of microorganisms involved in MIC, as well as understand their interactions. These limitations underscore the need for more robust and innovative approaches to studying MIC using more realistic laboratory simulations of offshore environments.

Schematic 1 provides an illustration of the proposed corrosion mechanisms for both the abiotic and biotic conditions during the initial stages, as they evolved over time during this present study.

**Schematic 1.**
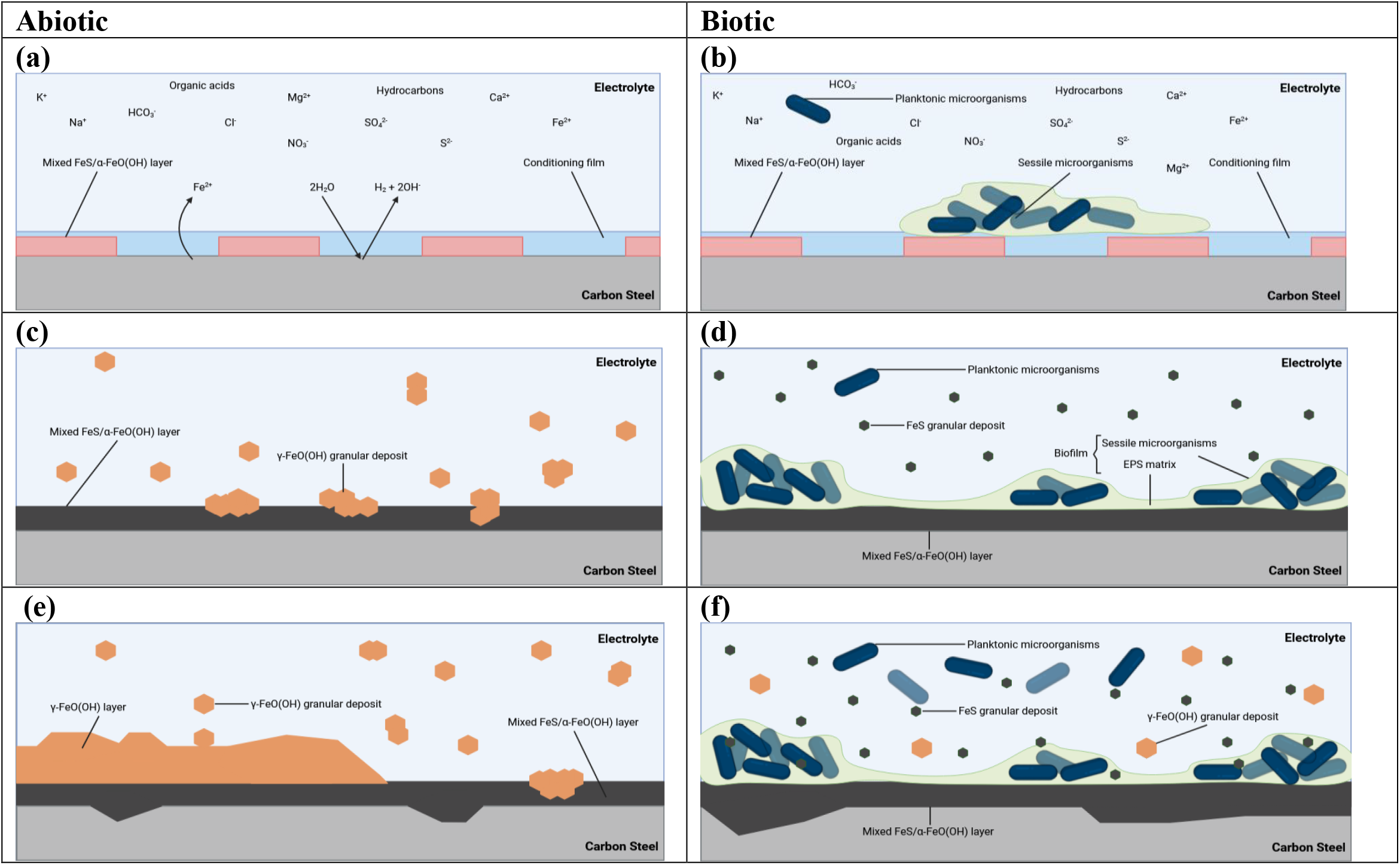
Illusrtation of the initial stages for UNS G10180 carbon steel in anaerobic abiotic and biotic PW media. corrosion mechanisms, (a, b) the formation of nascent inorganic corrosion film and the organic conditioning film with pioneering bacterial attachment during the initial batch phase; (c, d) maturing corrosion film under the abiotic condition with reduced biofilm growth and colonisation under the biotic condition due to the limited availability of organic carbon; (e, f) moderately low uniform and pitting corrosion under patchy corrosion deposits and thin biofilm with increasing granular deposits. BioRender.com (2024).

### Abiotic reactor

**-** After the initial batch phase, there was a general electronegative shift associated with anodic polarisation for the abiotic condition. A pseudo-steady state *E*_corr_ had not been attained after 28 days. PW often contains various organic compounds, including hydrocarbons and organic acids [37]. Additionally, PW typically contains inorganic compounds such as salts, minerals, and metal ions. Both organic and inorganic substances can precipitate on metal surfaces, contributing to the formation of a conditioning film. This film can affect the electrochemical properties of the steel surface, potentially making it more susceptible to corrosion or providing passivation. From the EIS ECMs, there is initially a capacitive behaviour observed over the first week, which reflects ion adsorption and the development of a conditioning film (mixed organic and inorganic interfacial layer). Subsequently, there was a diffusive behaviour during the final three weeks in the film layer. Moreover, there was a diffusive behaviour in the metallic interface double layer, which reflects charge transfer, due to the formation of corrosion products. The primary corrosion product identified was mackinawite. There were also additional bands which may be attributed to sulphur, as well as reference iron oxide compounds such as magnetite, goethite, lepidocrocite and hematite. Additional analysis identified that corroded areas were mainly covered by Fe and O, with heterogeneous distribution of S. Generally, there was a moderately low level of uniform corrosion of the steel surface for the abiotic media.

### Biotic reactor

**-** During the initial batch phase, where additional organics were available via supplementation with yeast extract in the pre-culture, an electronegative shift in the *E*_corr_ was observed. This can be attributed to the increased organics and/or biofilm formation on the CS, as the PW became dark green/black in colouration with visible black precipitates and increased turbidity. However, once the flow of fresh media was initiated, there was a similar electronegative shift which was observed for the abiotic condition. The potentials for both abiotic and biotic conditions in the latter stages were generally similar. Additionally, the *R*_p_ was generally low for both conditions. For the biotic condition, there is initially a capacitive behaviour observed over the first week, with a diffusive behaviour over the final three weeks in the film layer. This was similar to the abiotic condition. Whereas there was a diffusive behaviour over the initial two weeks, followed by a capacitive behaviour over the final two weeks in the double layer. This capacitive behaviour may be attributed to a biofilm, which may cause diffusion limitations. Likewise, the formation of corrosion products, which were primarily identified as mackinawite, will also be impacting the possible electrochemical reactions that are taking place at the interface of the metal/electrolyte.

Surface profilometry analysis provided further insights and revealed that there were low levels of uniform or localised pitting corrosion present for both the abiotic and biotic condition. However, after 28 days, the biotic condition did exhibit pits with a greater average area. This is characteristic of localised pitting caused by biofilms [38, 39]. Whilst localised pitting corrosion did not appear to be significantly exacerbated by the biotic condition, longer-term studies may reveal critical stages of biofilm maturation and corrosion progression. For this study, it was not possible to quantitatively determine *PD* values, due to the general absence of pitting across the coupon surfaces.

Analysis of the community dynamics revealed a marked change in the predominant relative abundances of microorganisms. The dominant genera from the sediment sample were generally anaerobic, halophilic, and obligately chemolithoautotrophic, obtaining energy by oxidizing inorganic compounds. Interestingly, the relative total archaea from the Day 0 planktonic sample still accounted for approximately 20% of the relative abundance. *Methanococcoides*, and *Methanococcus*, are both known methanogenic archaea that play an important role in the production of methane in anaerobic environments [40, 41]. The primary metabolic pathway in *Methanococcus* involves the reduction of CO_2_ with H_2_ to form methane [40]. Whereas *Methanococcoides* utilise methylotrophic methanogenesis, where methylated compounds serve as the primary substrates for methane production [41]. *Vibrio* was the dominant genera in these samples at approximately 30% relative abundance and are typically found in marine and estuarine environments. *Vibrio* sp. are known for their ability to form biofilms on various surfaces, including metals [42]. After 28 Days, there was a significant increase in *Marinobacter, Pseudomonas*, and *Halomonas*, making up approximately 80% of the relative abundance. *Marinobacter* species are known for forming biofilms on metal surfaces in marine environments. These biofilms can influence the corrosion process by altering the local chemical environment [43]. Moreover, *Marinobacter* species can degrade hydrocarbons, making them important for the bioremediation of oil spills in marine environments. They are well-adapted to high-salinity environments and play a role in iron cycling in marine environments, impacting the availability of iron [44, 45]. *Pseudomonas* species are prolific biofilm formers, and these biofilms can enhance corrosion by creating microenvironments that promote differential aeration and localized corrosion [46]. *Halomonas* species thrive in high-salinity environments and can reduce 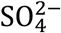 to H_2_S. Additionally, *Halomonas* species participate in nitrogen cycling, including denitrification, which is important for maintaining the balance of nitrogenous compounds in marine ecosystems [47]. The AR biofilm sample also had relatively high abundances of Proteobacteria species, namely *Marinobacterium* which made-up approximately 40%. Generally, the dominant species were halophilic or halotolerant and are known to be heterotrophic. They are well-adapted to marine conditions, playing roles in organic matter degradation, nutrient cycling, and interactions with marine organisms. Though the specific role of *Marinobacterium* and *Malaciobacter* are less characterised compared to other bacteria like *Pseudomonas* or *Halomonas*, they occupy a similar phenotypic niche [48]. Interestingly, there was an increase in the relative abundance of *Shewanella* within the AR biofilm. *Shewanella* species are known for their unique role in MIC, primarily due to their ability to reduce metal ions and interact directly with metal surfaces. In the presence of other bacteria, *Shewanella* can enhance corrosion through indirect mechanisms, such as H_2_S production and the formation of FeS. Moreover, *Shewanella* have a demonstrated ability for EET [49], which is an important process in MIC. It would be interesting to conduct longer-term studies with the same experimental setup to observe the threat of *Shewanella*.

**Schematic 2.**
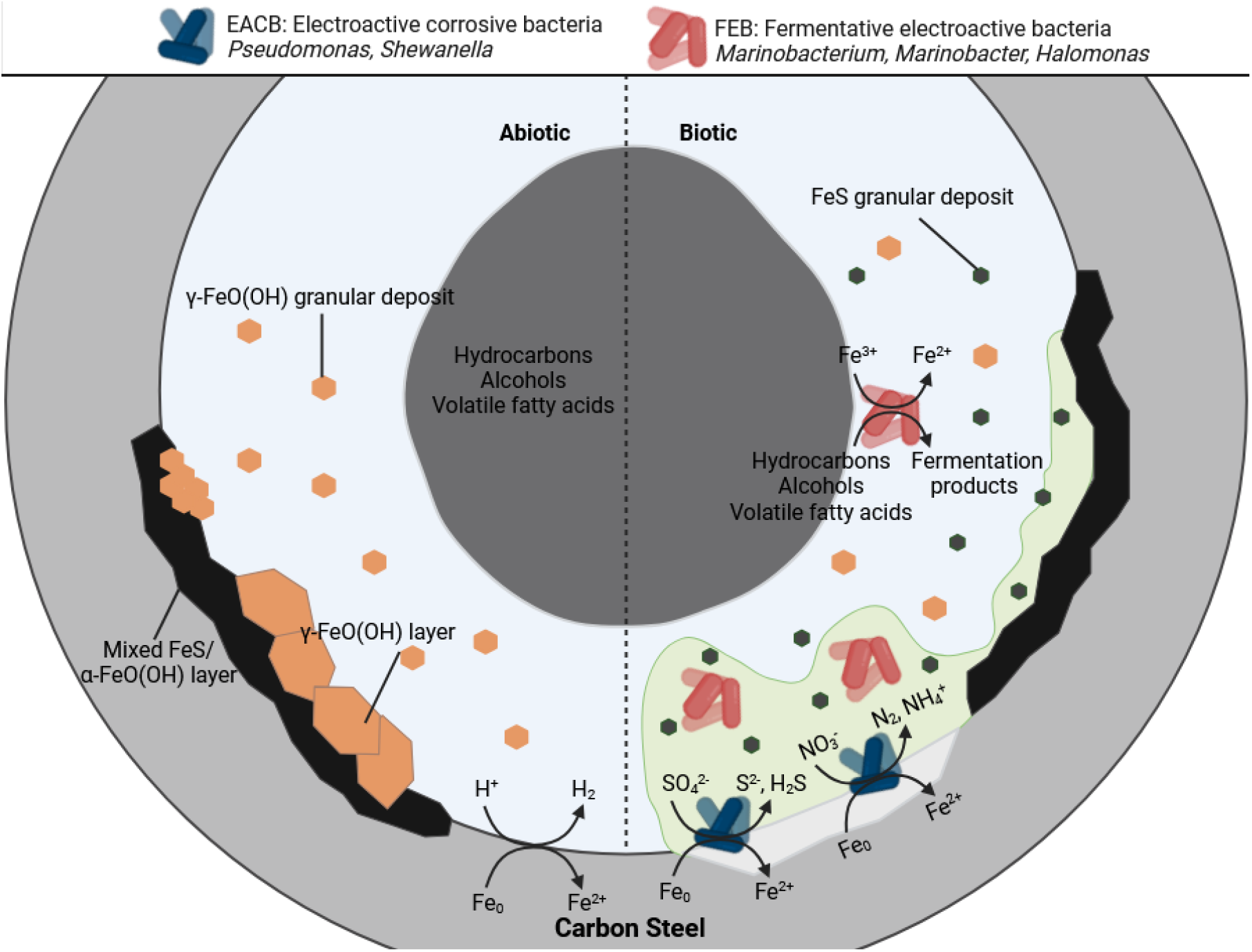
Overview of key reactions for UNS G10180 carbon steel in anaerobic abiotic and biotic PW media. It is important to note that the reactions illustrated may be complementary or antagonistic processes, particularly within the heterogeneous biofilm.

For the conditions used in this study, more closely mimicking the environmental conditions of an offshore oilfield system, the abiotic surfaces had the presence of a black corrosion product across the entire coupon surfaces with some reddish-brown granular deposits. As stated earlier, the primary corrosion product identified was mackinawite; with reference iron oxide compounds also detected. Mackinawite can form under anoxic or reducing conditions in environments containing H_2_S. H_2_S can be present in natural waters due to the reduction of 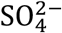. Moreover, mackinawite formation is favoured in anaerobic environments. Conversely, iron oxides are typically formed in the presence of O_2_ and water through oxidation reactions. Several types of iron oxides can form depending on environmental conditions. Initial electrochemical reactions produce iron hydroxides which are subsequently oxidised further to iron oxides. Nonetheless, there appeared to be a passivation of the metal surface as *CR*s were moderately low.

For this study, the electrochemical response under biotic conditions was not too dissimilar to the abiotic condition. However, surface observations and analysis of the biofilm suggest a different scenario. The biotic surfaces were only partially covered by a black corrosion product, with the presence of a heterogeneous dark green/black biofilm. As stated earlier, the mixed-species biofilm contained *Marinobacterium, Malaciobacter, Pseudomonas*, and *Halomonas* which all contribute to biofilm formation in *high salinity* marine environments. They are important players in nutrient cycling within marine ecosystems and are crucial in creating microenvironments. *In* mixed microbial communities, these bacteria can work synergistically by altering environmental conditions [44, 45, 46, 47]. Through the production of organics and other metabolites, that can influence the redox potential or pH, they can create microenvironments which may support 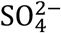 and Fe^0^ reduction over long-term studies. It is hypothesised that the mixed-species biofilm is acting as a diffusion barrier and providing some passivation of the metal surface. Schematic 2 proposes possible abiotic and biotic reaction mechanisms, associated primarily with the formation of mackinawite.

Previous laboratory studies have generally found that environmental conditions that mimic offshore oilfield systems, such as salinity, temperature, nutrient levels, and fluctuating O_2_ concentrations, have a profound impact on the formation, structure, and threat of corrosion from mixed-species biofilm communities on CS [50]. These studies emphasise the importance of replicating real-world offshore conditions in laboratory experiments to accurately assess and mitigate the threat of MIC in oil and gas systems. However, like *in situ* studies, there are limitations and challenges when it comes to designing laboratory studies. One of the primary challenges in these studies is accurately replicating the complex and dynamic environmental conditions found in offshore systems, such as fluctuating temperatures, pressures, and chemical compositions. Laboratory settings often simplify these conditions, which can limit the relevance of the findings to real-world applications. Many studies are constrained by time, making it difficult to observe long-term biofilm development and the full extent of MIC. Corrosion is a slow process, and short-term studies may miss critical stages of biofilm maturation and corrosion progression. Maintaining a stable and representative microbial community in laboratory settings is also challenging. In industrial environments, microbial communities are constantly evolving, and this dynamic nature is difficult to capture in a controlled environment. Laboratory studies often struggle to replicate the scale and flow conditions of actual pipelines. The differences in flow dynamics can significantly impact biofilm formation and corrosion patterns, meaning lab results may not fully translate to field conditions. Thus, designing experiments that closely mimic real-world conditions as much as possible is critical. Furthermore, long-term studies are needed to fully understand the impacts of biofilm communities on MIC in oil and gas systems.

## 5 Conclusions

In conclusion, by utilizing a marine sediment microbial consortium and replicating the environmental conditions of an offshore oilfield system within the laboratory, we gained valuable insights into biofilm development, community dynamics, and the potential for inducing MIC within a novel dual bioreactor protocol.

- The electrochemical response under both abiotic and biotic conditions was found to be similar. An initial conditioning film appeared to form, influencing the electrochemical properties of the steel surface. However, the PW in this study did not support rapid biofilm growth, likely due to the absence of rich carbon sources such as amino acids, peptides, and sugars.
- Mackinawite was identified as the primary corrosion product. Additionally, there were bands that may correspond to sulphur and reference iron oxide compounds, including magnetite, goethite, lepidocrocite, and hematite, observed under both conditions.
- Both conditions exhibited a moderately low uniform *CR* and limited localized pitting. Consequently, it was challenging to quantitatively assess *PD* values due to the general lack of pitting across the coupon surfaces. However, the biotic condition did display pits with a larger average area, indicative of MIC.
- Sequencing-based biofilm characterisation revealed that *Marinobacterium, Malaciobacter, Pseudomonas*, and *Halomonas* played critical roles in biofilm formation under conditions simulating an offshore oilfield system. These microorganisms are essential in nutrient cycling within marine ecosystems and contribute to the creation of microenvironments.

This study begins to bridge the gap between experimental findings and real-world scenarios involving mixed-species biofilms and MIC. The innovative dual bioreactor protocol, which leverages MLOE, enables a comprehensive understanding of initial biofilm formation and the metabolic changes occurring within the biofilm over time. Identifying and characterizing specific microorganisms under simulated environmental conditions is crucial for understanding the threat posed by MIC. If not detected and mitigated early, the microbial mechanisms can lead to significant costs. Ultimately, the aim is to enhance the effectiveness of biofilm management strategies in the industry, thereby improving sustainability.

## Data Availability

The raw/processed data required to reproduce these findings can be shared upon request.

## CRediT authorship contribution statement

**Liam Jones:** Conceptualization, Data curation, Formal Analysis, Investigation, Methodology, Project Administration, Validation, Visualization, Writing – original draft, Writing – review & editing. **Maria Salta:** Conceptualization, Writing – review & editing, Supervision, Project administration, Funding acquisition. **Torben Lund Skovhus**: Conceptualization, Writing – review & editing, Supervision, Project administration, Funding acquisition. **Kathryn Thomas:** Conceptualization, Writing – review & editing, Supervision, Project administration, Funding acquisition. **Timothy Illson:** Conceptualization, Writing – review & editing, Supervision, Project administration, Funding acquisition. **Julian Wharton:** Conceptualization, Writing – review & editing, Supervision, Project administration, Funding acquisition. **Jeremy Webb:** Conceptualization, Writing – review & editing, Supervision, Project administration, Funding acquisition.

## Acknowledgements

Many thanks to Dr Joe Parker who generously provided his expertise and assistance when it came to bioinformatics analysis. We are also grateful to Dr Terence Harvey and Dr Mark Willet for their assistances with Infinite Focus Microscopy and Confocal Laser Scanning Microscopy. Thanks to Dr Niall Hanrahan for his knowledge and guidance with Raman spectroscopy. Also, we are very grateful for the support from our colleagues in the Department at the University of Southampton. Finally, thank you to the Euro-MIC CA20130 - European MIC Network – New paths for science, sustainability and standard network for all the inspiration.

